# Balancing the Local and the Universal in Maintaining Ethical Access to a Genomics Biobank

**DOI:** 10.1101/157024

**Authors:** Catherine Heeney, Shona M. Kerr

## Abstract

**Background:** Issues of balancing data accessibility with ethical considerations and governance of a genomics research biobank, Generation Scotland, are explored within the evolving policy landscape of the past ten years. During this time data sharing and open data access have become increasingly important topics in biomedical research. Decisions around data access are influenced by local arrangements for governance and practices such as linkage to health records, and the global through policies for biobanking and the sharing of data with large-scale biomedical research data resources and consortia.

**Methods:** We use a literature review of policy relevant documents which apply to the conduct of biobanks in two areas: support for open access and the protection of data subjects and researchers managing a bioresource. We present examples of decision making within a biobank based upon observations of the Generation Scotland Access Committee. We reflect upon how the drive towards open access raises ethical dilemmas for established biorepositories containing data and samples from human subjects.

**Results:** Despite much discussion in science policy literature about standardisation, the contextual aspects of biobanking are often overlooked. Using our engagement with GS we demonstrate the importance of local arrangements in the creation of a responsive ethical approach to biorepository governance. We argue that governance decisions regarding access to the biobank are intertwined with considerations about maintenance and viability at the local level. We show that in addition to the focus upon ever more universal and standardised practices, the local expertise gained in the management of such repositories must be supported.

**Conclusions:** A commitment to open access in genomics research has found almost universal backing in science and health policy circles, but repositories of data and samples from human subjects may have to operate under managed access, to protect privacy, align with participant consent and ensure that the resource can be managed in a sustainable way. Data access committees need to be reflexive and flexible, to cope with changing technology and opportunities and threats from the wider data sharing environment. To understand these interactions also involves nurturing what is particular about the biobank in its local context.

## Background

In policy and governance discussions around the sharing of genetic and genomic data, the rights of research subjects are often balanced against the scientific benefits of allowing open access to biomedical data [1, 2]. Here, we follow an established tradition of including empirical worked examples, in order to engage with ethical issues raised by providing access to data stored within an existing biobank [3],[4],[5]. The aim is to go beyond a simple opposition between being open and protection of the autonomy and privacy of data subjects to make a case for the inclusion of social and technical considerations in assessing what is ethical. We seek to address a gap in discussion in scientific, law and policy arenas about standardisation [6],[7] by focusing on the contextual aspects of biobanking, which include the will and ability to sustain the resource [8]. We will consider Generation Scotland (GS) [9] as one context in which the global policy agenda of open access meets local issues. This raises not only questions around ethics and governance, which have been a focus of much of the discussion around access, but also questions of sustainability. GS is a genomics research biobank initiated with a Scottish Higher Education Funding Council grant between 2001 and 2004 and supported by the Chief Scientist Office of the Scottish Government from 2005 to 2014 (http://www.generationscotland.org). Participant recruitment and collection of data from the 24,000 plus participants began in 2006 and a GS infrastructure continues to exist and manage access to a repository more than a decade later. At the time of its creation Generation Scotland was described as: a “large, family-based intensively-phenotyped cohort recruited from the general population across Scotland, as a resource for studying the genetics of health areas of current and projected public health importance” [10].

The period of the early 2000s saw genetic biobanks and repositories of various sorts created with the aim of ensuring a greater openness to data sharing. Generally, biobanks or repositories hold both genetic data and phenotypic data sourced from research clinics and often also eHealth records. UK Biobank and Generation Scotland were intended to be resources with minimal restrictions to reuse [8]. Data and sample repositories such as GS were a manifestation of a growing commitment coming from science policy actors, including funders, to promote data sharing and access to a wide range of users. In what follows, we consider the specific characteristics of the GS repository in shaping its data access practices in light of the UK data sharing policy environment. We will use our interaction with GS to explore in particular the issues of sustainability and evolution of the content in relation to ethical decision making around access.

## Methods

We engage with GS as a specific example of a biobank and our “encounters with experience” [4] gained within the Generation Scotland Access Committee (GSAC). Empirical studies of the perspectives of those running biobanks point to the need to consider the contextual aspects of biobanking [11],[12],[13]. We hope to further elucidate these contextual aspects by focusing on the practices of the GSAC. The authors have both had experience of working within the GSAC. Using GS as a case enables us to consider how a particular repository attempts to balance locally established governance and institutional and research relationships with the imperative to share data as openly as possible. Through GS we explore the entwined practical and ethical challenges around data sharing for existing repositories [14]. By situating the ethics of access via examples arising in an active biobank, the aim is to ensure that our discussion goes beyond considerations of the “what if” type, which for example balance future health benefits against potential privacy risk questions for an imagined future [15].

We reflect upon the processes within GS through which requests for data access are handled. This will include considering how the Access Committee must deal creatively and responsively with issues not foreseen when the repository was first set up and as a consequence, which may challenge existing governance arrangements [16]. Therefore, we present examples of the decision-making process of the GSAC in order to consider how changes in the global data sharing and governance environment, as well as internal changes to the resource, raise ethical questions. It is in the context of the GSAC that the wider policy field, is negotiated in relation to the characteristics of GS. Developments within the biobank and the data sharing environment more generally can raise ethical dilemmas even if the consent obtained was comparatively broad. ‘While in an unproblematised situation, material objects are seamlessly woven into every day practice - family pedigrees are produced, blood samples are taken, medical records are filed away etc. In some cases the movement of these material objects into different physical locations, or even the particularity of their arrangement or configuration, can serve to problematize those practices’ [17]. We argue that data sharing raises issues relating to the role of the repository in future governance and ethical oversight and how expectations and preferences of participants will be interpreted in future scenarios, for example within consortia [18]. Whilst issues of sustainability could be viewed as separate from ethical and governance considerations, we aim to show that they are inseparable in relation to access [19].

## Results

### Generation Scotland (GS)

The GS Scottish Family Health Study was designed to provide a research resource, adequately powered to detect moderate sized genetic effects upon common and chronic disease and traits. A family-based recruitment strategy was employed to collect over 24,000 participants between 2006 and 2011. Study participants were first approached through their General Practitioner (GP) using the Community Health Index (CHI) number (the CHI number exists only in Scotland and is unique to each individual in the >96% of the Scottish population registered with a GP) [20]. Those who indicated that they and one or more of their relatives would participate were sent an information leaflet, a consent form and a preclinical questionnaire. Subsequently, comprehensive information was collected covering demographics, biometric measurements and the health of individuals and their families, including psychological health. This was done via a paper questionnaire which gathered ∼400 data items, and during a research clinic appointment a further ∼150 items of data were collected. The membership of the GSAC includes representation from NHS Research and Development Offices, University Technology Transfer Offices, clinical academics, scholars working on ethics and governance, and laboratory and IT experts. The membership is renewed over time but individuals connected with the research design, recruitment and maintenance have ongoing input. A dedicated management group responsible for implementing the access arrangements was funded by the Chief Scientist Office (CSO) of the Scottish Government, for an initial period of three years, following the completion of recruitment (the management structure of GS is illustrated in Figure 1). Currently, members of the GS Executive Committee, Expert Working Groups and Access Committee give their time freely as part of their academic or support staff duties.

**Figure 1.**
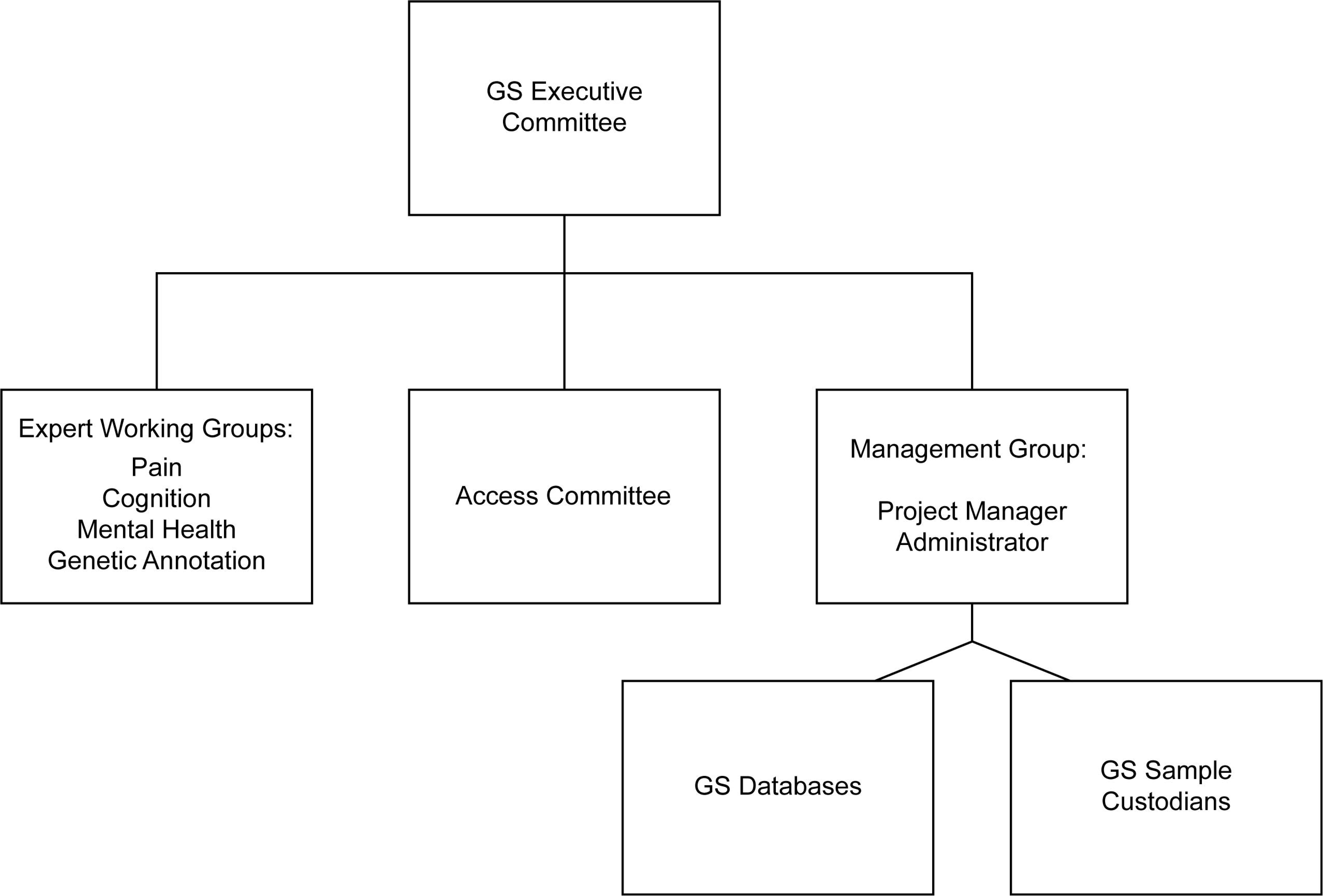
Generation Scotland (GS) Management Structure

The Management, Access and Publication Policy of GS specifies that the Access Committee (GSAC) will: manage requests for collaboration and use of Project Data, Derived Data and NHS Data and/or Samples; Approve or deny requests for new collaborations; Consider and approve: Collaboration Proposal Forms and Data and Material Transfer Agreements and Report to the Executive on the progress of proposed collaborations. It must also consider the terms of consent and protection of confidentiality of data subjects [9]. GSAC meetings are held approximately quarterly depending on the volume and complexity of the proposals. The GSAC considers a range of criteria when reviewing data access requests, as is standard practice for Data Access Committees operating a managed (controlled) access process [21],[22]. Pre-screening by the GS Management Group ensures that full reviews take place only after funding has been obtained by the applicants and if requested resources match GS holdings. Requests which are considered routine are dealt with via email without detailed discussion by the GSAC. Routine requests seek to access data only, do not require the participants to be re-contacted and are viewed as raising no significant governance issues. Examples include the re-analysis of existing anonymised genomic datasets to test and improve software algorithms, or to access anonymised individual patient data according to a particular genotype of interest and to generate preliminary data for a grant application.

Furthermore, GSAC is notified but is not directly involved in decisions to approve release of GS data and/or samples for management rather than research purposes to the GS management team or to academic researchers from institutions that are part of the GS Collaboration Agreement. Approximately 20 projects out of a total of over 250 research requests have required this type of release, which are not subject to Data and Material Transfer Agreement signature or an access charge. The primary purpose of these management access requests is to test or check the quality of an aspect of the resource. In contrast, payment must be made for all research requests, as it was decided by the GS Executive Committee that no distinction would be made between researchers who had been involved in the GS Collaboration Agreement and those who had not.

### Policy, access and repositories in the UK

Influential genomics and medical research funder the Wellcome Trust has long advocated data sharing and open access [8]. In 2003 the Wellcome Trust published an influential report, following a closed meeting in Fort Lauderdale, Florida, intended to identify and resolve issues that may form barriers to data sharing and reuse of existing biomedical data [23]. Here a model of access was promoted in which any studies with biomedical research value could become a “community resources” [23], in opposition to what were presented as restrictive and proprietorial governance models. This meeting itself followed on from a meeting in Bermuda held in February 1996, which involved leaders of the Human Genome Project and sought to avoid some scientists gaining an advantage by refusing to share human sequence information in a timely manner [24]. More recently a meeting was held in Toronto in part to deal with technological advances in data production. This produced the Toronto Statement of 2009, which sought to reiterate the importance of reaffirming the ‘lessons from Human Genome Project (HGP)’ regarding the ‘scientific value’ of rapid release of data [25]. In these statements, collective benefits of data sharing are often in counter-balance to more individual goods such as privacy and professional protectionism [26]. This generates tensions between notions of public and societal benefit from open access to large and complex biomedical datasets and the need to manage access with respect to the details of participant consent and privacy expectations [27],[28]. The Toronto Statement acknowledges that under some circumstances, such as where detailed genomic or clinical data pose a risk of deidentification of individual research subjects, ‘access may be restricted’ [25]. Some of the policy documents discussed in this section, as we will point out, do acknowledge privacy and confidentiality issues raised by data-sharing and access. Moreover, the linked issues of privacy and confidentiality have been raised as a concern across literature produced by stakeholders in biomedical data sharing [29]. Further disadvantages put forward include ‘moral distance’ between new contexts for data use and sharing and compromising the ability of original researchers to fulfill commitments to research subjects. Questions about adequate acknowledgement of those involved in maintaining the study were also raised [8],[29]. The governance response for extending access has often been upon altering models of consent [30],[31]. Existing resources such as the Avon Longitudinal Study of Parents and Children and the 1958 British Birth Cohort were encouraged to revisit their participant consent [32]. Newer large scale repositories such as Generation Scotland and UK Biobank, created to support research access by secondary users [30], attempted to begin with broad informed consent to facilitate more flexible data access arrangements.

The policy environment for data sharing continues to abound with examples of encouragement to share. For example, data deposition to repositories such as the European Genome-phenome Archive (EGA) [33], a service for permanent archiving and sharing of all types of personally identifiable genetic and phenotypic data resulting from biomedical research projects, is included as a condition of publication for many key scientific journals. However, access to resources remains frustratingly difficult for some, prompting suggestions that access arrangements for biobanks tread a fine line between facilitating and hindering sharing [34]. More recently the Wellcome Trust, with the Expert Advisory Group on Data Access (EAGDA), has produced a report addressing incentives and disincentives to sharing for funders, institutions and researchers [35]. Again it is acknowledged in this document that restrictions can be mandated ethically due to confidentiality concerns for study participant data. However, the EAGDA report emphasises the importance of recognising the benefits of data sharing, including avoiding duplication of effort and allowing innovative and inventive uses of data already collected [35]. A Concordat produced by Research Councils UK (RCUK) [36], which is an umbrella organization for the main public sector research funding bodies in the UK, echoed these benefits and added the potential for safeguarding against misconduct. The RCUK Concordat does not deal exclusively with data arising from human subjects, and is thus largely silent on the matter of informed consent and the management of ongoing relationships with research participants. However, Principle 5 deals with the need for some elements of managed access. Justified restrictions to data access include the protection of commercial interests and again the privacy and confidentiality of research subjects. These documents aim for generic guidance in which strategies for dealing with these issues appear as high level principles. Whilst there is acknowledgement of justified ethical restrictions to sharing, the onus is placed on those controlling data sharing to make the case for denying access [36]. A joint review of data security, consent and opt-outs in the UK National Health Service (NHS) and Care Quality Commission in 2016 saw Dame Fiona Caldicott, the National Data Guardian, re-emphasise the responsibility of organisations using NHS data to ensure both anonymity and consent [37]. Yet there is an increasingly warm view of sharing health related data in the 2012 Caldicott report [38], where a seventh principle added to the list of six produced in 1997 states that sharing patient data could be viewed as equally important as the protection of patient confidentiality. Indeed, access to NHS data has been promoted to meet both healthcare and commercial interests [38].

The EAGDA suggests that consent ‘is not a panacea and even where consent specifically allows for further data use, robust governance is essential for the ethical conduct of research’ suggesting a forbearing attitude to managed access approaches [35]. The report devotes considerable space to ensuring that governance and other institutional arrangements promote data sharing where possible. This report does then engage with issues which may arise in particular contexts but opts to list a generic set of challenges, largely focused on concerns around protecting the privacy of individuals.

The uniqueness of the genome of an individual coupled with even minimal phenotypic trait information are widely accepted to pose risks to data confidentiality and by extension the privacy of individual data subjects [28],[39],[40]. Controversy around the publication of a key paper in 2008 [41], which highlighted the possibility of re-identifying an individual within an anonymized dataset [42] led to reactive policy changes around data access by key players, such as the Wellcome Trust and the National Iinstute for Health in the U:S [1],[8]. Confidentiality and security arrangements and use restrictions are intended to mitigate against such incidents, which are thought may have the indirect effect of undermining trust in a particular repository, or in biomedical research as a whole [28]. The EAGDA has acknowledged the difficulties of maintaining promises of anonymity, suggesting that misuse including attempts at identification of individual data subjects will potentially incur both legal and funding sanctions [35]. Meanwhile those who have sought to promote open and rapid data sharing have shown a tendency to frame privacy concerns as a barrier to research. Privacy and related issues such as confidentiality and anonymity have themselves become a target for those who are convinced of the benefits of open access, with the suggestion that actual harms arising, even where some breach is possible, are exaggerated [43].

The UK policy environment reflects an international effort to take practical steps to ensure existing data is shared by improving the discoverability of biobanks, such as the UK MRC Cohort Directory (http://www.mrc.ac.uk/research/facilities-and-resources-for-researchers/cohort-directory/) and the UK CRC Tissue Directory (http://www.biobankinguk.org/). The latter is part of an umbrella organization for biobanking in Europe, BBMRI-ERIC (Biobanking and Biomolecular Resources Research Infrastructure), funded by the European Commission [44]. GS is listed in these directories and also tries to make its resources findable through the established academic routes of a study website and research publications as well as exploiting social media channels.

Recently, an international initiative for scholarly data publishing proposed that all scientific data should be "FAIR"-Findable, Accessible, Interoperable, and Reusable [45]. The Global Alliance for Genomics and Health (GA4GH) has developed a Framework for responsible sharing of genomic and health-related data [46]. Here the push is towards creating the conditions of more open sharing via harmonisation: “The Global Alliance is working to alter the current reality where data are kept and studied in silos, and tools and methods are non-standardized and incompatible” (http://genomicsandhealth.org/). The hope is that a more standardised approach to “tools and methods” used for collection, storage and characterisation of data will engender further improvements in the ease of data sharing.

### GS: where open access imperatives and local infrastructure meet

Whilst the emphasis coming from the wider science and data sharing policy environment encourages the prioritisation of data sharing [47], translating this into practice remains a local enterprise. Requirements for managing access to resources held in biobanks and biorepositories are inevitably interpreted at the level of projects, repositories or institutions [34]. The entwined discourse of standardisation and open access would suggest that such heterogeneity is detrimental. However, scholarship in the field of science and technology studies suggests that despite efforts at standardization, practice will necessarily maintain some aspects of a given context [48]. Unlike for example UK Biobank, GS has made a commitment to oversee and manage overlap in the research goals of applicants. Where it is likely that two separate applications would overlap, a situation that has arisen no more than 10 times in the history of GS, leading to a potential duplication of effort, GS offers to put the researchers in touch with the “Expert Working Group” (EWG) leads (Figure 1) [http://www.ed.ac.uk/generation-scotland/about/management/expert-working-groups].

However, there is no requirement for the applicant to collaborate with an EWG. Academics closely associated with the resource as part of the Expert Working Groups and the GS Executive Committee (Figure 1) continue to invest their time and intellectual capital in GS. Some of these academics were involved in the scientific design of the study, the recruitment of participants and convincing funders of its merit. Moreover, they continue to be involved in what has been termed ‘articulation work’ [49]. That is to say they (and others) have sought research funding for a variety of studies via which they have added further data to the resource, helping to maintain its relevance and scientific importance.

The EWGs include high profile academics whose expertise covers a particular area of research. Whilst a decision on the part of an applicant not to collaborate with the EWGs does not necessarily create a barrier to access, the EWGs constitute part of a commitment by GSAC to manage project overlap. A distinctive feature of access arrangements for GS relates to co-authorship, which is stated in the data and materials transfer agreement and the GS Authorship & Acknowledgement Policy [9]. GS requires that collaboration with a research group requesting access leads to shared authorship of research publications resulting from use of the GS resource. Whilst there are few sanctions that GS can apply to ensure compliance with this requirement for attribution of credit, there is just one example of these terms not being honoured from over 150 completed research projects with more than 100 published research papers. Some of the experts in governance suggest that the independence of access decisions form these sorts of contextual factors is desirable [50]. However, this is arguably a way in which the researchers and the repository can ensure reward and continued ethical and scientific input are in place. Principle 7 of the RCUK Concordat recognises the costs to research teams involved in data sharing, emphasising the importance of finding appropriate ways of acknowledging and rewarding those who collect and manage the data [36].

Requests for access to data only or data plus samples are submitted via a secure online portal, where researchers will also indicate whether linkage to NHS records or participant re-contact will be necessary. Four areas of evaluation (scientific, governance, data and materials) are then completed by designated GSAC members. The scientific assessment addresses questions around the methods and scientific contribution of the proposal. The governance assessment will often attempt to balance ethical considerations such as confidentiality guarantees and existing participant consents with details of the requested access to the resource and participants. Issues such as whether the participants will need to be re-contacted and on what basis are dealt with here. On one occasion, a proposal was declined as it asked for specific phenotypic information which was thought likely to raise sensitivities such as participants feeling that they had been “singled out”. The data assessment is usually done by a member of the GS management group who also sits on the GSAC. This will consider the data holdings of GS in respect of the type of data requested, flagging up practical issues relating to release of data including the relevant participant consent for linkage of their GS data and medical records via the CHI number for re-contact, or sharing samples outside the UK. For example, if an applicant has requested the samples to be sent overseas or has asked to re-contact participants, the data assessment will include a report on the number of individuals who have consented to this. On another occasion a proposal requested unusually specific data relating to only a small number of individuals and was returned with a request for an overhaul of the research design, on the grounds that it could pose a potential re-identification risk. The materials assessment is carried out when samples as well as data are requested. This assessment must consider whether the proposed research is an appropriate use of the physical samples, which are a finite resource. Projects looking simultaneously at multiple biomarkers in a high proportion of the cohort are preferred to those which measure only one biomarker in a small subset, as this creates more new data for the quantity of sample used. The proposal form also includes sections on why the GS resource was chosen to carry out the research and what benefits could be expected to accrue to GS as a result of providing access. Applicants are asked about whether ethical approval has been sought from their own institution, funder or other appropriate body. How to ensure appropriate recognition of work done in creating and maintaining the biobank in regard to each proposal is also a question that is raised depending on the type and extent of access sought [8]. Discussions during the GSAC meetings are in most cases attempts to accommodate these various aspects of the proposal alongside the question of sustainability of the resource. Issues can be and often are resolved by asking applicants for further information or modification of the type and scope of access sought in line with the concerns raised in the assessments, which are then discussed in the GSAC meetings.

### Access and sustainability

GS access arrangements and the discussion conducted as part of the GSAC meetings aim to strike a balance that promotes the sustainability of the resource whilst making it a ‘community resource’ [23]. One of the recommendations made by the EAGDA [35] is that repositories should be well-resourced, presumably to support the sort of activities that comprise the managed access approach undertaken by GS. Following the end of the period of funding from the Scottish Government, the GS project manager and administrator posts have been underwritten by a mixture of cost recovery charges for access and financial support from an NHS Research & Development fund. Cost recovery through access fees payable by researchers wishing to access the GS resource is a key part of sustainability, helping to maintain the lean institutional structure necessary to the governance and curation of the GS resource. It also provides an incentive to facilitate access, mitigating against any possible tendency towards withholding data [50]. Different tiers of pricing are in place for access to data and samples by academic and commercial applicants. As cost recovery via access fees is an important component in the sustainability of the study, it has been necessary to strike a balance between meeting the real overhead costs of maintaining GS and keeping charges at non-prohibitive levels. A steadier source of income would undoubtedly lead to a recalibration of current arrangements, which lend another layer to the decision making process around allowing access. At the time of writing, only four projects have not gone ahead due to an unwillingness by prospective secondary users to pay an access charge.

Sustainability questions also arise in relation to the wider data sharing environment and the existence of cost free alternatives to accessing genotype and phenotype data [19] broadly similar to that in GS. For example, access can be sought to resources via routine academic collaboration with the Principal Investigator of different cohorts, or from genomics data resources such as the EGA [33]. Another evolving aspect of the data environment impacting upon the GS biobank is requests for data to be released so that it can be housed on platforms elsewhere. This would diminish the position of GSAC as a single gatekeeper of the data. A current example can be seen in the collation and sharing of cohort data via the MRC Dementias Platform UK (http://www.dementiasplatform.uk/). Moves to release data into scientific databases that are publically accessible or have less or perhaps more restrictive modes of permitting access raise uncertainties around ethics and governance potentially creating ‘tremendous consent challenges’ [51].

The move in human genomics research towards collaborative working in very large international consortia, as exemplified by the Cohorts for Heart and Aging Research in Genomic Epidemiology (CHARGE) Consortium (http://www.chargeconsortium.com/) raise challenges in terms of recognition of effort. GS is one of the cohorts contributing data to CHARGE (and many other national and international genomics consortia). These consortia facilitate genome-wide association study meta-analyses and replication opportunities among multiple large and well-phenotyped longitudinal cohort studies. Data from GS contributes towards greater statistical power for new discoveries to be made and leads to co-authorship of members of the GS Executive on the resulting research papers. However, because data and summary statistics from a large number of cohorts are combined in a meta-analysis, there is a considerable distance between the details and nuances of the governance of each individual cohort and the data analytical research activities, such as producing summary statistics, within the consortium. For example, data analysts working in a consortium will usually not have been involved in the data access application made to each repository. This disconnect is evidenced by the resulting research papers often being co-authored by hundreds of researchers, with the description of each cohort usually confined to the online supplementary information. It also again raises potential ethical questions around expectations embedded in relationships between study participants and data producers [29],[32].

Attempts to mitigate these sorts of concerns can be seen in to the central genomics database dbGaP, where emphasis has been place upon the independence of the Data Access Committees (DACs) of this resource [50]. Independence in this scenario is interpreted as ‘without the involvement of data producers’ [50]. Other examples of this sort of centralised structure for managing access can also be found in the UK. For example, the Wellcome Trust Sanger Institute DAC manages access to data assembled for the Wellcome Trust Case Control Consortium and other projects located there [52]. While there is an increasingly acknowledged need to attribute credit for the role played by repositories such as GS, these efforts remain partial and aspirational in terms of setting standards [50]. Moreover, these efforts again appear to lean toward harmonisation to facilitate a decontextualisation of the data. However, context can be important both in terms of the significant time invested by researchers in the creation and maintenance of the repository and in regard to relationships with the study participants [8],[16]. Questions of how to incorporate the specifics of local practices, which respond to both sustainability issues and the expectations of research subjects within these large multi-repository, multi-institutional arrangements, are not directly addressed by ensuring the independence of Data Access Committees for repositories with a centralised governance system.

### Managing access to an evolving resource

One of the problems of attempting to produce enduring standardised access procedures is that unlike Access Committees, written protocols cannot respond to relevant developments both within and outside of individual biobanks. The emphasis place upon informed consent can be seen as problematic in this regard [16]. In the decade since GS began recruitment there have been changes both in the wider data sharing environment and in the composition of the repository itself. Written consent was sought and gathered during the original GS recruitment phase (from 2006) for study data to be linked to the NHS health records of participants, using their CHI number. This identifying number is used for all NHS Scotland procedures (registrations, attendances, samples, prescribing and investigations) and allows healthcare records for individuals to be linked across time and location [20]. Ethical approval for the record linkage was obtained (as part of the GS:SFHS Research Tissue Bank Approval) from the East of Scotland Research Ethics Committee. In each record linkage project, permissions were obtained by researchers from the NHS Privacy Advisory Committee or its successor, the Public Benefit and Privacy Panel for Health and Social Care, for use of NHS medical data.

The number of events, and measurements recorded, increase over time as the participants get older, which means there is an enormous additional pool of research relevant data about GS participants obtained *since* recruitment. Although initial data collection was cross-sectional, GS became a prospective cohort as a result of the ability to link to routine NHS data [53].

This makes the GS resource valuable for a new generation of researchers with evolving methodical and technological tools. In large part due to the existence of the Scottish CHI number and participant consent, GS is able to link to NHS records, wherein the scale of the data is vast. For example, in the biochemistry dataset alone, there are more than two million test results for over 800 measures relating to 11,000 GS participants in NHS Tayside, going back over 25 years. In a recent genomics research project using the GS resource, outputs from just over two thousand participants for one of these biochemical measures, uric acid, were tested for association with over 24 million genetic markers in a genome-wide association study (GWAS) [54]. Such research, employing new techniques including genotype data imputation, continues to augment the number of data items held on individual study participants within the repository^i^.

### GS and Evolving Participant Consent

Alongside anonymisation of data, informed consent is a standard tool for ensuring data access remains ethical in the face of such dynamic developments. It was acknowledged from the outset that it would be difficult to predict the precise nature of use of the GS resource in the future. For that reason, and due to the logistics and potential confidentiality challenges of contacting and re-consenting individual participants for each use, broad consent was sought from participants. This was intended to permit a very wide range of potential biomedical research (including commercial) uses^ii^. These documents have been subject to minor updates to reflect changes in the project. The latest versions (from early 2010) contain information relating to access to the resource and the management and protection of participant data. It is made clear that “Any access will be subject to the strictest ethical scrutiny and scientific rigour’ (GS PIL 2010). Generation Scotland participants originally consented to their data being made available to researchers from any sector worldwide. However, this original consent did not specifically allow samples to leave the UK, so additional consent for samples to be sent abroad was later obtained for a little under half the cohort (Table 1).

**Table 1.** Summary of consents in the GS resource at baseline (2006 – 2011) and added subsequent to the end of participant recruitment in 2011

| **Dataset** | **Date in GS** | **Participant Numbers** |
| --- | --- | --- |
| Participants recruited | 2006-2011 | 24,084 |
| Participants with consent and mechanism for record linkage | 2006-2011 | 22,014 |
| Participants with consent and mechanism for recontact | 2006-2011 | 21,992* |
| Participants with consent for sample transfer ex-UK | 2012-2013 | 11,255 |
| Participants with consent for recontact by email | 2016 | 6,546 |

\*This figure includes 785 participants known to have died since participation in GS

This additional consent was achieved via a re-contact exercise carried out in 2012-3 in response to several requests to allow the materials and data to be analysed using technologies only available in other countries. Re-contact for consent, as with all re-contact with study participants, was done through an established mechanism that, again, used the CHI number, with letters sent by post by an NHS intermediary for confidentiality. GS saw this as the most appropriate means to ensure scientifically and ethically valid research was not declined on the basis of the location of the laboratories. The additional consent process required a submission to a Research Ethics Committee and took well over a year from initiation to conclusion. It came at a considerable cost in terms of GS staff time and the non-negligible cost of postage to and from 21,207 individuals (88% of the 24,084 people in the study database, excluding participants who had died or who had not given consent for re-contact). Just over half this number replied with 11,255 participants giving consent for their samples to leave the UK (53% of people contacted in total). The decision to reconsent was a response to unforeseen changes in the scientific environment and clearly illustrates the reality of the challenges to relying upon consent as a unique means of ensuring that changing access practices are rendered ethical [31]. Although policy makers dealing with the protection of health data are taking an increasingly permissive view about the relationship between consent and access to medical records [38], GS continues to employ a consent based approach for use of these data. The joint report from the UK National Data Guardian was clear on the responsibility of organisations using NHS data to ensure not only appropriate consent but also anonymity of disseminated data [37]. Due to the detailed and potentially identifiable nature of the data collected in GS, it was agreed from the outset that the data would be released using a managed access policy [9]. From a technical point of view the duty to protect participant data from identification is dealt with by GS via IT and access arrangements. GS study participants’ information is protected by personal identifiers being held separately from all other study data, using an encrypted version of the CHI number. The encryption key is held within the NHS IT network and cannot be directly accessed by GS. Researchers as part of their host institution have to sign a Data User Agreement before they are allowed to access NHS-linked data. Linked eHealth data is released for clinical academic research via NHS approved safe havens, or held in a secure network environment [55]. Samples are stored in four separate laboratories in Scottish University Medical Schools, catalogued through a central laboratory information management system [56]. Only the GS management team holds the key between the sample identifiers and the phenotype data. These measures are designed to ensure that GS samples, which include DNA, blood, serum and urine, are not accessed without the appropriate authorisation via the access processes detailed above. Knowledge of these systems for protecting the data, the data subjects and reputation of the repository is part of what members of the GSAC bring to decisions about data release.

## Discussion

One of the issues highlighted in the paper is the interaction between specific local characteristics of a given biobank or repository, especially in relation to governance and sustainability, and the guidelines and ideals pervading the wider data sharing and science policy and ethics environment, which aim at harmonisation and fewer barriers to access [19]. The aim is to show how this policy context translates into the ways in which data is accessed and ethics and governance are enacted, given factors local to the repository and its practices. One part of this is the ability to respond to what is non-routine, as well as what was not foreseen when consent arrangements were entered into originally [17]. The GSAC in its decision-making must consider and accommodate a number of issues, in which it is difficult to make a clean separation between questions of sustainability, governance and ethics.

GSAC routinely discusses issues relating to the welfare of the data subjects, how further recontacts may be unduly onerous and whether or not a particular request for new data may make the data subject question their health status. Such questions relate to the continued goodwill of participants and raise sustainability considerations as do sharing arrangements which do not adequately recognise the academic and administrative work involved in providing GS data. Commitment of academics involved with the resource through the EWGs, the Executive Committee and the GSAC is an important part of ensuring that the governance model agreed to by participants is maintained.

In considering requests for access, GSAC must “ensure the Project, through its collaborations, conforms to the consent and ethical approval obtained, is not brought into disrepute and that participant confidentiality is respected” (Generation Scotland Management, Access and Publications Policy 2016). The work on ethics done by GSAC and the Management Group is interrogating what is being proposed to ensure continued alignment between the governance framework, participant expectations, the ability to manage the resource and strong encouragement from the scientific community to be as open as possible [35]. The first research proposal was approved by GSAC in 2008 (at a time when participant recruitment and data collection were ongoing) and the first research findings resulting from an access request were published in 2010. Nearly a decade after its formation, GSAC remains actively involved in mediating imperatives to promote access and ethical and sustainable research and management of the GS resource. We suggest that in addition to the focus upon ever more universal and standardised practices, the local expertise gained in the management of such repositories must also be nurtured and encouraged [48].

## Conclusions

In summary, a commitment to open access in genomics research has found substantial backing in science and health policy circles in the UK and beyond, as evidenced by the stance taken by research funders and the outputs of influential meetings such as those held in Bermuda, Fort Lauderdale and Toronto. This is not to stay that stakeholders are unaware of or unsympathetic to the issues around data subject privacy and to a lesser extent the interests of the data-producing scientists and institutions (both of which are mentioned in the Toronto Statement). However, in this paper we have attempted to throw light on how decisions on access sustainability can figure in solutions to potential problems for both data-producing scientists and research participants. Repositories of data and samples from human subjects may have to operate under managed access, to protect privacy, align with the participant consent and ensure that the resource can be managed in a responsive and sustainable way. Research on best-practice which looked at literature produced by stakeholders also found that some of the questions we have raised here, such as continuing to balance the interests of different stakeholders, fulfilling commitments to research participants and the ability to control analyses that may cause reputational damage, were ranked among the disadvantages of data sharing [29]. We have used our own engagement with GS in order to construct an argument about the importance of considering the local aspects to responsively accommodate access policies designed for universal application. Data access committees need to be reflexive and flexible, to cope with changing technology and opportunities and threats from the wider data sharing environment. These considerations are particularly relevant in relation to closure of a biobank [19],[57] an event that raises practical issues such as transfer of data or materials to other entities [58]. We have aimed to show that the responsive ethics work done by GSAC and counterparts in other smaller repositories is key in mediating between the global and the local [58] in the era of consortium working. Whilst ever greater emphasis is placed upon open access to data as a commercial and economic good, some mechanism for incorporating the role now carried out by the access committee will remain necessary.

## Declarations

### Ethics approval and consent to participate

The processes of participant recruitment to GS:SFHS received ethical approval from the NHS Tayside Committee on Medical Research Ethics (REC Reference Number: 05/S1401/89).

GS:SFHS has subsequently been granted Research Tissue Bank status by the Tayside Committee on Medical Research Ethics (REC Reference Number: 15/ES/0040), providing approval for a wide range of uses within medical research, including genetic analyses and eHealth record linkage.

## Consent for publication

Not applicable

## Availability of data and material

The data supporting this article are in the public domain.

## Authors’ contributions

Both authors contributed to the writing of the manuscript, in an iterative manner. The text was drafted by CH and SK, who have each read and approved the final manuscript.

## Acknowledgements

We thank David Porteous, Reka Nagy, Shawn Harmon and Gill Haddow for constructive comments on drafts of the manuscript. We are indebted to the GS Project Manager and GS Administrator, Archie Campbell and Laura Boekel, for their helpful responses to our requests for information on details of access requests and issues arising from these.

## Footnotes

i 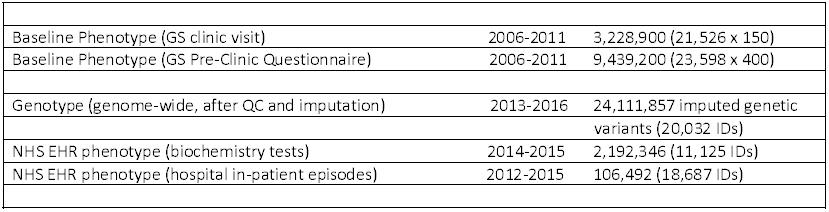

ii Examples of consent forms and Participant Information Leaflets can be viewed on the GS website (www.generationscotland.org).

## References

1. Craig DW, Goor RM, Wang Z, Paschall J, Ostell J, Feolo M, Sherry ST, Manolio TA: Assessing and managing risk when sharing aggregate genetic variant data. Nat Rev Genet 2011, 12:730–736.

2. Laurie G: Reflexive governance in biobanking: on the value of policy led approaches and the need to recognise the limits of law. Hum Genet 2011, 130:347–356.

3. Haimes E, Williams R: Sociology, ethics, and the priority of the particular: learning from a case study of genetic deliberations. Br J Sociol 2007, 58:457–476.

4. Ives J: ‘Encounters with experience’: empirical bioethics and the future. Health Care Anal 2008, 16:1–6.

5. Dunn M, Sheehan M, Hope T, Parker M: Toward methodological innovation in empirical ethics research. Camb Q Healthc Ethics 2012, 21:466–480.

6. Kaye J: Do we need a uniform regulatory system for biobanks across Europe? Eur J Hum Genet 2006, 14:245–248.

7. Ouellette S, Tasse AM: P(3)G - 10 years of toolbuilding: From the population biobank to the clinic. Appl Transl Genom 2014, 3:36–40.

8. Kaye J, Heeney C, Hawkins N, de Vries J, Boddington P: Data sharing in genomics--re-shaping scientific practice. Nat Rev Genet 2009, 10:331–335.

9. http://www.ed.ac.uk/generation-scotland/using-resources/access-to-resources/access-policy: Generation Scotland. Accessed 15/03/2017.

10. Smith BH, Campbell H, Blackwood D, Connell J, Connor M, Deary IJ, Dominiczak AF, Fitzpatrick B, Ford I, Jackson C, et al: Generation Scotland: the Scottish Family Health Study; a new resource for researching genes and heritability. BMC Med Genet 2006, 7:74.

11. Kaye J GSMC, Heeney C, Smart A, Parker M, (Eds): Governing Biobanks: Understanding the Interplay Between Law and Practice. Hart Oxford 2012.

12. Cadigan RJ, Lassiter D, Haldeman K, Conlon I, Reavely E, Henderson GE: Neglected ethical issues in biobank management: Results from a U.S. study. Life Sci Soc Policy 2013, 9:1.

13. Muddyman D, Smee C, Griffin H, Kaye J: Implementing a successful data-management framework: the UK10K managed access model. Genome Med 2013, 5:100.

14. Murtagh MJ, Turner A, Minion JT, Fay M, Burton PR: International Data Sharing in Practice: New Technologies Meet Old Governance. Biopreserv Biobank 2016, 14:231–240.

15. Hanson VL: Envisioning Ethical Nanotechnology: The Rhetorical Role of Visions in Postponing Societal and Ethical Implications Research. Science as Culture 2011, 20:1–36.

16. Heeney C: An “Ethical Moment” in Data Sharing. Sci Technol Human Values 2017, 42:3–28.

17. Parker M: Ethnography/ethics. Social Science & Medicine 2007, 65:2248–2259.

18. Kaye J, Hawkins N: Data sharing policy design for consortia: challenges for sustainability. Genome Med 2014, 6:4.

19. Chalmers D, Nicol D, Kaye J, Bell J, Campbell AV, Ho CW, Kato K, Minari J, Ho CH, Mitchell C, et al: Has the biobank bubble burst? Withstanding the challenges for sustainable biobanking in the digital era. BMC Med Ethics 2016, 17:39.

20. Pavis S, Morris AD: Unleashing the power of administrative health data: the Scottish model. Public Health Res Pract 2015, 25:e2541541.

21. Dyke SO, Kirby E, Shabani M, Thorogood A, Kato K, Knoppers BM: Registered access: a ‘Triple-A’ approach. Eur J Hum Genet 2016, 24:1676–1680.

22. Shabani M, Knoppers BM, Borry P: From the principles of genomic data sharing to the practices of data access committees. EMBO Mol Med 2015, 7:507–509.

23. Sharing Data from Large-scale Biological Research Projects: A System of Tripartite Responsibility [https://wellcome.ac.uk/sites/default/files/wtd003207_0.pdf]

24. Marshall E: Bermuda rules: community spirit, with teeth. Science 2001, 291:1192.

25. Toronto International Data Release Workshop A, Birney E, Hudson TJ, Green ED, Gunter C, Eddy S, Rogers J, Harris JR, Ehrlich SD, Apweiler R, et al: Prepublication data sharing. Nature 2009, 461:168–170.

26. Lowrance W: Learning from experience: privacy and the secondary use of data in health research. Research report. Nuffield Trust. Research report Nuffield Trust 2002.

27. Sheehan M: Can Broad Consent be Informed Consent? Public Health Ethics 2011, 4:226–235.

28. Heeney C, Hawkins N, de Vries J, Boddington P, Kaye J: Assessing the privacy risks of data sharing in genomics. Public Health Genomics 2011, 14:17–25.

29. Bull S, Roberts N, Parker M: Views of Ethical Best Practices in Sharing Individual-Level Data From Medical and Public Health Research: A Systematic Scoping Review. J Empir Res Hum Res Ethics 2015, 10:225–238.

30. Cambon-Thomsen A: Science and society - The social and ethical issues of post-genomic human biobanks. Nature Reviews Genetics 2004, 5:866–873.

31. Wallace SE, Gourna EG, Laurie G, Shoush O, Wright J: Respecting Autonomy Over Time: Policy and Empirical Evidence on Re-Consent in Longitudinal Biomedical Research. Bioethics 2016, 30:210–217.

32. Heeney C and Smart A: Enacting Governance - The Case of Access. Governing Biobanks: Understanding the Interplay between Law and Practice 2012:232–258.

33. Lappalainen I, Almeida-King J, Kumanduri V, Senf A, Spalding JD, ur-Rehman S, Saunders G, Kandasamy J, Caccamo M, Leinonen R, et al: The European Genome-phenome Archive of human data consented for biomedical research. Nat Genet 2015, 47:692–695.

34. Fortin S, Pathmasiri S, Grintuch R, Deschenes M: ’Access arrangements’ for biobanks: a fine line between facilitating and hindering collaboration. Public Health Genomics 2011, 14:104–114.

35. https://wellcome.ac.uk/sites/default/files/governance-of-data-access-eagda-jun15.pdf: Establishing Incentives and Changing Cultures to support Data Access. 2015.

36. Concordat on Open Research Data [http://www.rcuk.ac.uk/documents/documents/concordatonopenresearchdata-pdf>]

37. Guardian ND: Review of Data Security,Consent and Opt-Outs https://www.gov.uk/government/uploads/system/uploads/attachment_data/file/535024/data-security-review.PDF. 2016.

38. The Information Governance Review: To Share or Not to Share DoH: 2012.

39. Erlich Y, Narayanan A: Routes for breaching and protecting genetic privacy. Nat Rev Genet 2014, 15:409–421.

40. Gymrek M, McGuire AL, Golan D, Halperin E, Erlich Y: Identifying personal genomes by surname inference. Science 2013, 339:321–324.

41. Homer N, Szelinger S, Redman M, Duggan D, Tembe W, Muehling J, Pearson JV, Stephan DA, Nelson SF, Craig DW: Resolving individuals contributing trace amounts of DNA to highly complex mixtures using high-density SNP genotyping microarrays. PLoS Genet 2008, 4:e1000167.

42. Clayton D: On inferring presence of an individual in a mixture: a Bayesian approach. Biostatistics 2010, 11:661–673.

43. Savage N: Privacy: The myth of anonymity. Nature 2016, 537:S70–72.

44. Mayrhofer MT, Holub P, Wutte A, Litton JE: BBMRI-ERIC: the novel gateway to biobanks. From humans to humans. Bundesgesundheitsblatt Gesundheitsforschung Gesundheitsschutz 2016, 59:379–384.

45. Wilkinson MD, Dumontier M, Aalbersberg IJ, Appleton G, Axton M, Baak A, Blomberg N, Boiten JW, da Silva Santos LB, Bourne PE, et al: The FAIR Guiding Principles for scientific data management and stewardship. Sci Data 2016, 3:160018.

46. Knoppers BM: Framework for responsible sharing of genomic and health-related data. Hugo J 2014, 8:3.

47. Bobrow M: Funders must encourage scientists to share. Nature 2015, 522:129.

48. Timmermans S, Berg M: Standardization in action: Achieving local universality through medical protocols. Social Studies of Science 1997, 27:273–305.

49. Fujimura JH: Constructing Do-Able Problems in Cancer-Research - Articulating Alignment. Social Studies of Science 1987, 17:257–293.

50. Shabani M, Dyke SO, Joly Y, Borry P: Controlled Access under Review: Improving the Governance of Genomic Data Access. PLoS Biol 2015, 13:e1002339.

51. Caulfield T, McGuire AL, Cho M, Buchanan JA, Burgess MM, Danilczyk U, Diaz CM, Fryer-Edwards K, Green SK, Hodosh MA, et al: Research ethics recommendations for whole-genome research: consensus statement. PLoS Biol 2008, 6:e73.

52. https://wellcome.ac.uk/sites/default/files/governance-of-data-access-annexes-eagdajun15.pdf: GOVERNANCE OF DATA ACCESS: ANNEXES. 2015.

53. Smith BH, Campbell A, Linksted P, Fitzpatrick B, Jackson C, Kerr SM, Deary IJ, Macintyre DJ, Campbell H, McGilchrist M, et al: Cohort Profile: Generation Scotland: Scottish Family Health Study (GS:SFHS). The study, its participants and their potential for genetic research on health and illness. Int J Epidemiol 2013, 42:689–700.

54. Nagy R, Boutin TS, Marten J, Huffman JE, Kerr SM, Campbell A, Evenden L, Gibson J, Amador C, Howard DM, et al: Exploration of haplotype research consortium imputation for genome-wide association studies in 20,032 Generation Scotland participants. Genome Med 2017, 9:23.

55. Lea NC, Nicholls J, Dobbs C, Sethi N, Cunningham J, Ainsworth J, Heaven M, Peacock T, Peacock A, Jones K, et al: Data Safe Havens and Trust: Toward a Common Understanding of Trusted Research Platforms for Governing Secure and Ethical Health Research. JMIR Med Inform 2016, 4:e22.

56. Macleod AK, Liewald DC, McGilchrist MM, Morris AD, Kerr SM, Porteous DJ: Some principles and practices of genetic biobanking studies. Eur Respir J 2009, 33:419–425.

57. Stephens N, Dimond R: Closure of a human tissue biobank: individual, institutional, and field expectations during cycles of promise and disappointment. New Genet Soc 2015, 34:417–436.

58. Zawati MH, Borry P, Howard HC: Closure of population biobanks and direct-to-consumer genetic testing companies. Hum Genet 2011, 130:425–432.

